# Simulation-based inference for close-kin mark-recapture: implications for small populations and nonrandom mating

**DOI:** 10.1101/2024.12.16.628779

**Authors:** Paul B. Conn

## Abstract

Close-kin mark-recapture (CKMR) uses data on the frequency of kin pair relationships (e.g. parent-offspring, half siblings) in genetic samples from animal populations to estimate parameters such as abundance and adult survival probability. To date, most applications of CKMR have relied on a pseudo-likelihood approximation where pairwise comparisons of relatedness are assumed to be independent. This approximation works well when abundance is high and the sampled fraction of the population is low (as with many marine fisheries), but resultant estimators have inflated nominal confidence interval coverage when applied to small populations. Small populations and nonrandom mating structures also lead to problems with using second-order kin for estimation because one cannot typically differentiate half-siblings from other kin pair types like aunt-niece. In this paper, I assess the utility of approximate Bayesian computation (ABC) as a way of relaxing the pseudo-likelihood independence assumption. Under this approach, one only needs the ability to simulate population and sampling dynamics and to summarize resulting statistics in an informative way (e.g., number of kin pairs of different types). After exploring bias and interval coverage in several simulation studies, I illustrate these procedures on CKMR data from a Canadian caribou population. I show that ABC substantially improves interval coverage, and allows inference for difficult biologies where it would be difficult to calculate the analytical probabilities necessary for a binomial pseudo-likelihood. Simulation-based approaches to inference have great potential for expanding the application of CKMR to small populations and for breeding dynamics that are difficult to model.

---

Close-kin mark-recapture (CKMR) uses data on the frequency of kin pair relationships (e.g. parent-offspring, half siblings) in genetic samples to make inference about population parameters such as abundance and adult survival (Skaug, 2001; Bravington et al., 2016). Unlike traditional capture-recapture, which requires longitudinal samples in which animals have the opportunity to be captured more than once, CKMR methods can potentially be applied to single sampling events such as fish and wildlife removals. Because knowledge of abundance and survival are fundamental to ecological research and animal conservation, there is now considerable interest in applying CKMR models to fish and wildlife populations. For instance, CKMR has been applied to commercial fisheries (Bravington, Grewe and Davies, 2016; Rawding, Sharpe and Blankenship, 2014; Wacker et al., 2021), freshwater fishes (Prystupa et al., 2021; Ruzzante et al., 2019), elasmobranchs of conservation concern (e.g. sharks, skates, rays; Hillary et al., 2018; Patterson et al., 2022; Trenkel et al., 2022), and marine and terrestrial mammals (Lloyd-Jones et al., 2023; Merriell, Manseau and Wilson, 2024; Taras et al., 2024). There are considerably more CKMR studies currently in progress or planned, so such applications will only continue to grow.

To provide intuition about how CKMR works, consider a simplified setup where there are *N*_*f*_ adult females and *N*_*o*_ offspring, and we desire to estimate the number of adult females in the population. Obtaining genetic samples from *n*_*f*_ adult females and *n*_*o*_ offspring and conducting a parentage analysis, we observe a binary random variable *Y*_*ij*_ for each pair that equals one if female *i* is the mother of *j* and zero otherwise. For instance, in Fig. 1, we have *N*_*f*_ = 8, *N*_*o*_ = 6, and *n*_*f*_ = *n*_*o*_ = 4. There are a few constraints to note about Fig. 1. First, each offspring has a single mother. Second, each female has a maximum of one offspring in this example, although this may vary in other species based on the biology of the species (litter size, etc.). Given that each offspring “marks” its mother, we can view this single sampling event as anolagous to a simple two sample mark-recapture experiment. In our example, there are *n*_*o*_ = 4 marks, *n*_*f*_ = 4 is the size of a second sample used to look for presence of marks, and *m* = 2 is the number of marks found in the second sample. A Lincoln-Petersen estimator (see e.g. Seber, 1982) is then 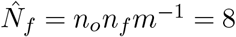 in our example (clearly, numbers were chosen judiciously).

**Fig 1.**
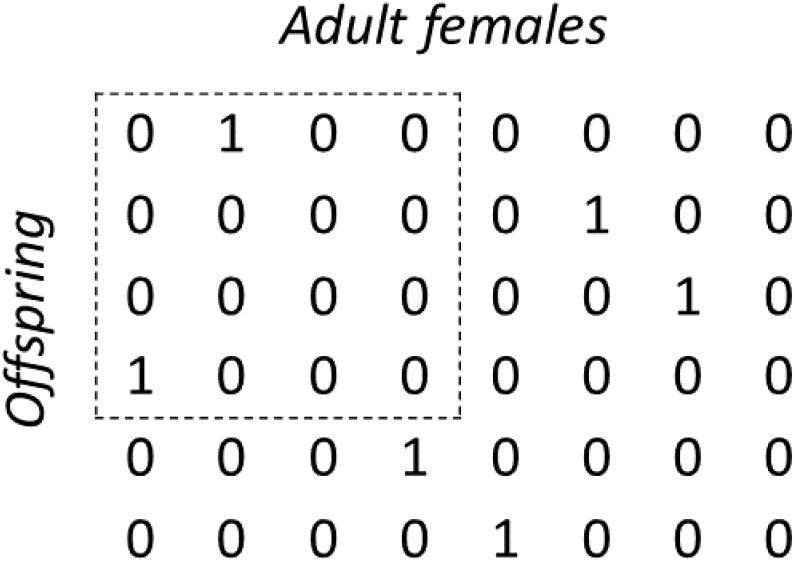
A simple, hypothetical population of six offspring and eight adult females. In this example, each offspring has a single mother, and each mother has a maximum of one offspring; close-kin inference focuses on using realized kinship relationships within a single sample of the four offspring and four adult females (dashed region) in order to conduct inference about adult female population size.

Although a few close-kin studies have employed Lincoln-Petersen-type estimators (Rawding, Sharpe and Blankenship, 2014; Ruzzante et al., 2019), it is much more common for close-kin studies to use pseudo-likelihood for statistical inference (Skaug, 2001; Bravington et al., 2016). Under this approach, one simply compares each genetic sample to every other one, treating each pairwise comparison as independent. This approximation works well for large populations and when a relatively small proportion of the population is sampled. It also lends itself more readily to model extensions, such as multiple years of sampling, inclusion of population trend parameters in the pseudo-likelihood, and consideration of other kin pair types (e.g., siblings). Within the single sample framework, however, it is easy to see the danger of applying pseudo-likelihood inference to data from small populations: one is essentially approximating a hypergeometric distribution with a binomial, which will lead to inflated variance estimates (Merriell, Manseau and Wilson, 2024). Consider again Fig. 1; a pseudo-likelihood in this instance would treat the state-space of possible observed kin-pairs (the dashed area) as consisting of 16 zeroes, 16 ones, and anything in between, effectively ignoring the constraints that rows and columns sum to one. Ideally, precision estimates would reflect such known constraints.

Another difficulty with CKMR inference is the confounding that can happen between different kin pair types. For instance, half-siblings, grandparent-grandchild, and full thiatic pairs (e.g., full aunt-niece) all have the same amount of co-inherited DNA so cannot usually be discriminated using genetics alone. Although it is possible to model such confounded data using a mixture of probabilities (Bradford et al., 2018; Taras et al., 2024), to our knowledge no one has yet written down a probability of a full thiatic pair within a CKMR model. Full thiatic pairs are unlikely in nature with large populations and randomized mating, but are much more likely when repeat breeding occurs between two animals over time, which can happen with small populations or with species that exhibit pair bonding. In such situations, CKMR inference has thus far been restricted to parent-offspring pairs.

Although it may be prohibitively difficult to write down full likelihoods associated with full pedigrees, it is comparatively easy to simulate them. For instance, armed with knowledge of vital rates (e.g., survival, fecundity schedules), mating dynamics, and spatial population structure, there are several software packages that allow researchers to simulate population dynamics and mating events and to keep track of kin pair types (e.g., Anderson, 2022; Baylis, 2019). The relative ease of simulating data suggests that simulation-based inference (i.e., approximate Bayesian computation (ABC); Beaumont, Zhang and Balding, 2002) may be a viable alternative to pseudo-likelihood inference. Indeed, ABC has a rich history in analysis of population genetics data where exact likelihoods border on intractable (e.g., Boitard et al., 2016).

In this paper, I explore the use of simulation-based inference to (1) more accurately characterize precision of CKMR estimators for small populations, and (2) to permit CKMR estimation with biologies and sampling scenarios that make analytical calculation of kinship probabilities difficult. I do this through a sequence of simulation studies, before analyzing a “real life” caribou data set.

## 1. Simulation studies

I start by considering a simple, one sample situation where a closed population estimator can be used to estimate animal abundance using data on the frequency of parent-offspring pairs (POPs). This scenario lets us compare alternative CKMR estimators when the distribution of the data is known exactly, both in terms of bias and variance. In a second scenario, I consider a multi-year study where abundance and exponential population trend are of interest, again using POPs. In a third scenario, I look at using CKMR to estimate abundance and survival from POPs, full sibling pairs (FSPs), and a confounded category of half-sibling pairs (HSPs), full thiatic pairs (FTPs) and grandparent-grandchild pairs (GGPs) when there is a monogamous mating structure. Although my focus is not on developing ABC algorithms per se, I consider increasingly sophisticated ABC algorithms through these examples, as necessitated by increasing dimensions of the parameter space and possible number of summary statistics. Throughout, I do not assume the reader has prior knowledge of ABC, but the reader may also wish to consult reviews (Beaumont, Zhang and Balding, 2002; Beaumont, 2010, e.g.) or online lectures (e.g. Filippi, 2019) to help develop intuition. To maintain continuity, I describe results of each simulation study immediately following the methods description.

### 1.1 Simulation study 1: Single sample

Consider a population that is sampled in a similar manner to Fig. 1; that is, a genetic sample of *n*_*f*_ adult females is obtained out of a population of *N*_*f*_, where *N*_*f*_ is unknown. Similarly, a sample of *n*_*o*_ offspring is obtained from a population of *N*_*o*_. Letting *y*_*ij*_ = 1 if individual *i* is *j*’s mother (and zero otherwise), we seek to estimate the population of mothers (i.e., *N*_*f*_). For instance, *N*_*f*_ could be the number of spawning female salmon in a river, and *N*_*o*_ could be the number of outmigrating smolt (Rawding, Sharpe and Blankenship, 2014).

#### 1.1.1. Data generation

To generate data from this scenario, I used a factorial design, involving two options for adult female abundance, *N*_*f*_ ∈ {100, 400}, and two types of offspring distributions, including: (i) all adult females have a single offspring, and (ii) all adult females have two offspring. For each combination of these factors, I generated 2000 datasets. For the *N*_*f*_ = 100 scenario, I set *n*_*f*_ = *n*_*o*_ = 50, and for the *N*_*f*_ = 400 scenario I set *n*_*f*_ = *n*_*o*_ = 100. In each case, individuals were sampled at random without replacement.

#### 1.1.2. Estimation models

In the specific case where each mother *i* produces a single offspring *j*, where no mortality occurs before sampling, and where sampling of the population is random with respect to genotype, it can be shown that the probability mass function for the number of sampled mother-offspring pairs 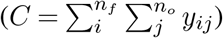 is

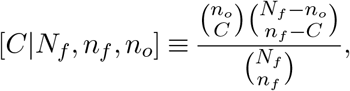

which is recognizable as a hypergeometric distribution. This formulation is not new, having provided the basis for the Lincoln-Petersen estimators in standard closed population capture-recapture analysis (Seber, 1982). Thus, we are in the rare case (for CKMR, anyway) of having a “true” likelihood for the data in at least one demographic scenario, estimation with which can be compared to other alternatives to assess their relative performance. However, note that the maximum likelihood estimator (MLE) for the hypergeometric index (*N*_*f*_) is known to be biased for small populations (Chapman, 1951).

In total, I fit six estimation models to each simulated dataset, including:

- The hypergeometric model, using maximum likelihood;
- A binomial model, using maximum likelihood (representing a standard CKMR pseudo-likelihood), where

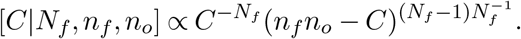 Note that the binomial pseudo-likelihood is motivated by comparing each offspring to each adult female (there are a total of *n*_*f*_ *n*_*o*_ such comparisons). Assuming independence among trials, the probability of a success is simply 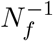;
- The Chapman estimator (Chapman, 1951), which includes a small population bias correction for the hypergeometric;
- Simulation-based inference using an “exact” approximate Bayesian Computation algorithm (“ABC-exact”);
- An inexact ABC algorithm in which parameter values associated with the the lowest 5% squared discrepancy values are retained (“ABC-quant05”);
- The ABC-quant05 scenario, but where local linear regression is used to improve performance (cf. Beaumont, Zhang and Balding, 2002) (“ABC-quant05-reg”).

For the first two estimation approaches, I used the MLE to assess bias and 90% profile likelihood intervals to summarize nominal confidence interval coverage. For the Chapman estimator, nominal confidence interval coverage was assessed relative to “0.5 transformed logit” intervals (Sadinle, 2009).

For the 3 ABC approaches to inference, I used Algorithm 1. To facilitate comparison with MLEs, I used the posterior mode as a point estimate (computed using a kernel density approximation), and computed 90% credible interval coverage relative to 0.05 and 0.95 quantiles. For the “ABC-quant05-reg” estimator, the credible interval calculation was performed using a weighted empirical cumulative distribution function (Baddeley, Rubak and Turner, 2015), constructed using the weights **W** from Algorithm 1.

Figure 2 shows profile likelihoods and approximate Bayesian posteriors (ABP) for 5 of the CKMR estimators applied to a single simulated data set with *N*_*f*_ = 100 and mothers having a single offspring. The ABP for the “exact” ABC method is almost identical to the hypergeometric profile likelihood, with the ABC-quant05-reg and ABC-quant05 estimators having slightly greater spread. The binomial profile likelihood has considerably greater spread than the other alternatives.

**Fig 2.**
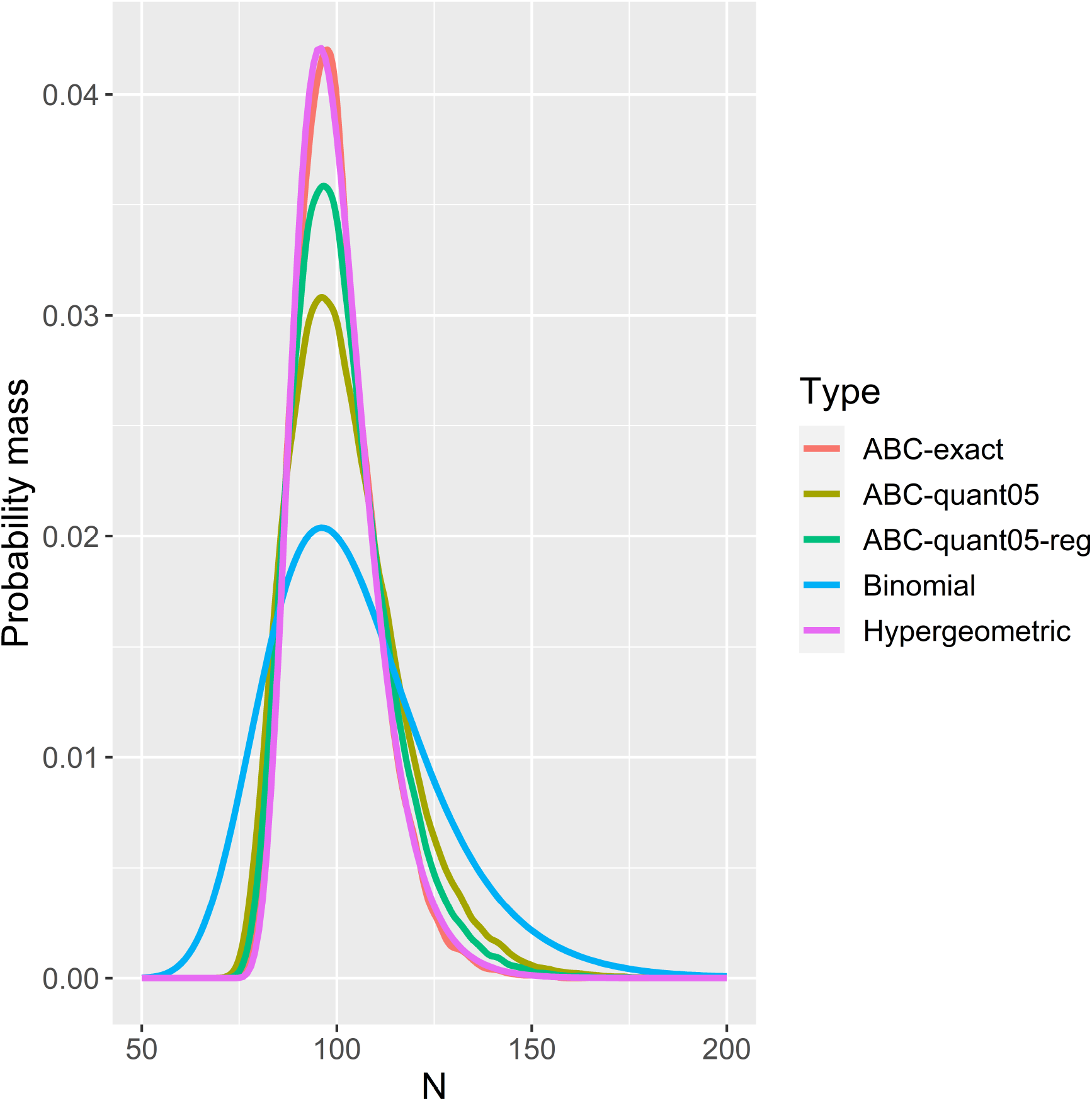
Likelihood profiles (binomial, hypergeometric) and posterior kernel density estimates (ABC methods) of adult female abundance resulting from analysis of a single simulated CKMR dataset. In this case, the hypergeometric is well approximated by the ABC-exact algorithm (which only retains samples where the number of simulated kin pairs is equal to the observed value). The other ABC algorithms based on the lowest 5% MSE discrepancy in kinship count have higher variance, but are closer to the true likelihood than the binomial “pseudo-likelihood.”

##### Algorithm 1

Approximate Bayesian computation algorithms for single sample CKMR data

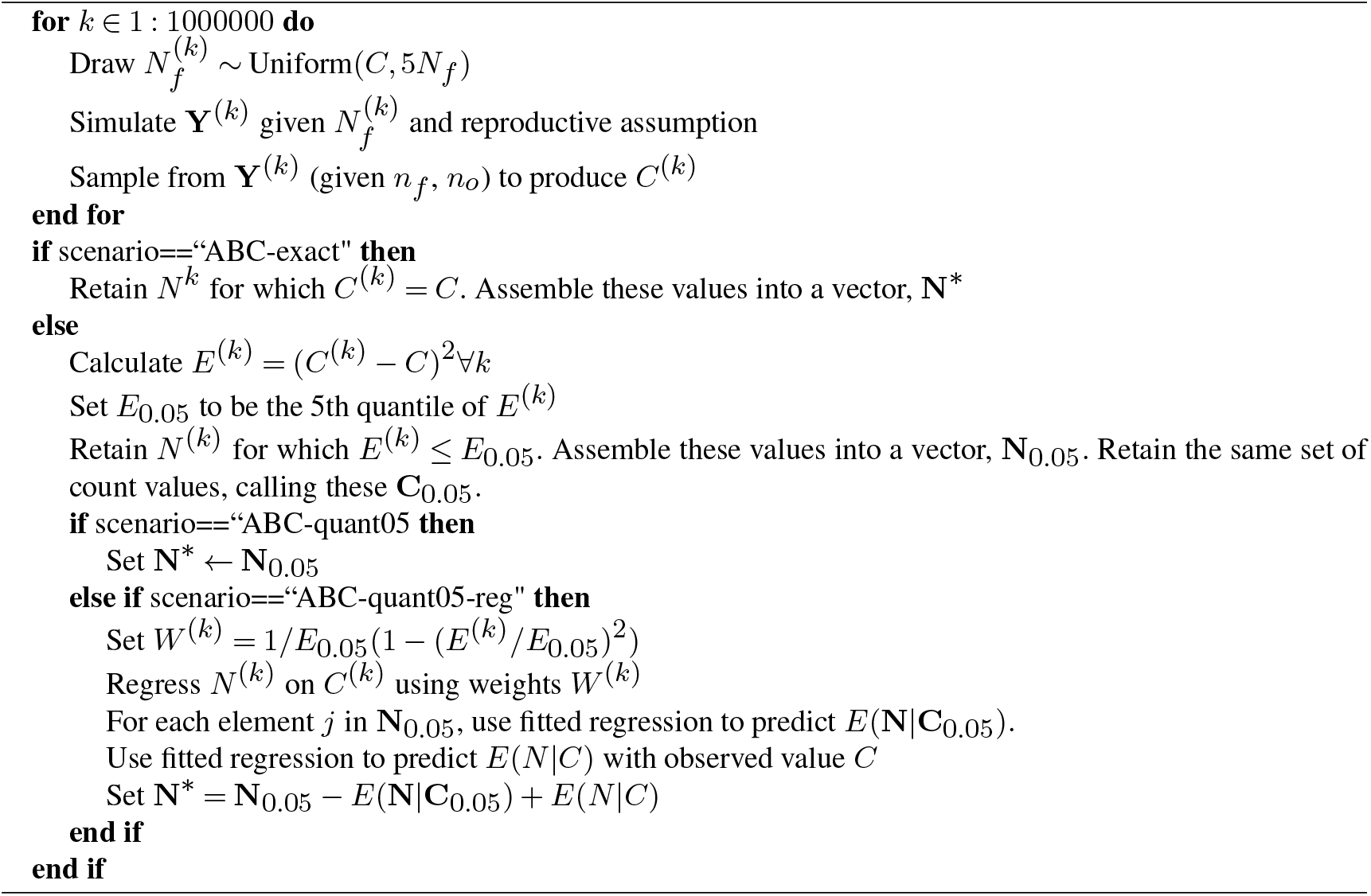

Over all the simulations, only the Chapman estimator appeared to be unbiased over all simulation inputs. The other five estimators had statistically equivalent positive biases within each of the scenarios, including:

- 1% for the *N*_*f*_ = 100, 1 offspring scenario
- 1.3% for the *N*_*f*_ = 100, 2 offspring scenario
- 2.5% for the *N*_*f*_ = 400, 1 offspring scenario
- 3.7% for the *N*_*f*_ = 400, 2 offspring scenario.

Although these biases were similar, estimators varied considerably when it came to nominal coverage (Fig. 3). In particular, the binomial estimator always overcovered, including close to 100% coverage for the *N*_*f*_ = 100, one offspring scenario, mirroring findings by Merriell, Manseau and Wilson (2024). The degree of overcoverage decreased with either more offspring per mother or greater adult female population size. The hypergeometric and Chapman estimators performed well when mothers had one offspring, but undercovered when mothers had two offspring, particularly for the *N*_*f*_ = 100 scenario. The ABC approaches tended to have coverage close to nominal, though the ABC-exact method tended to under-cover and the ABC-quant05 tended to slightly overcover. I suspect that some of the coverage results represent an interaction between interval length and observed positive bias; for instance, the interval length for the ABC-exact estimator is probably about right, but is shifted to the right by 1-3.5%. Similarly, the ABC-quant05-reg scenario likely has interval lengths that are slightly too long, but are sufficient to compensate for the observed positive bias.

**Fig 3.**
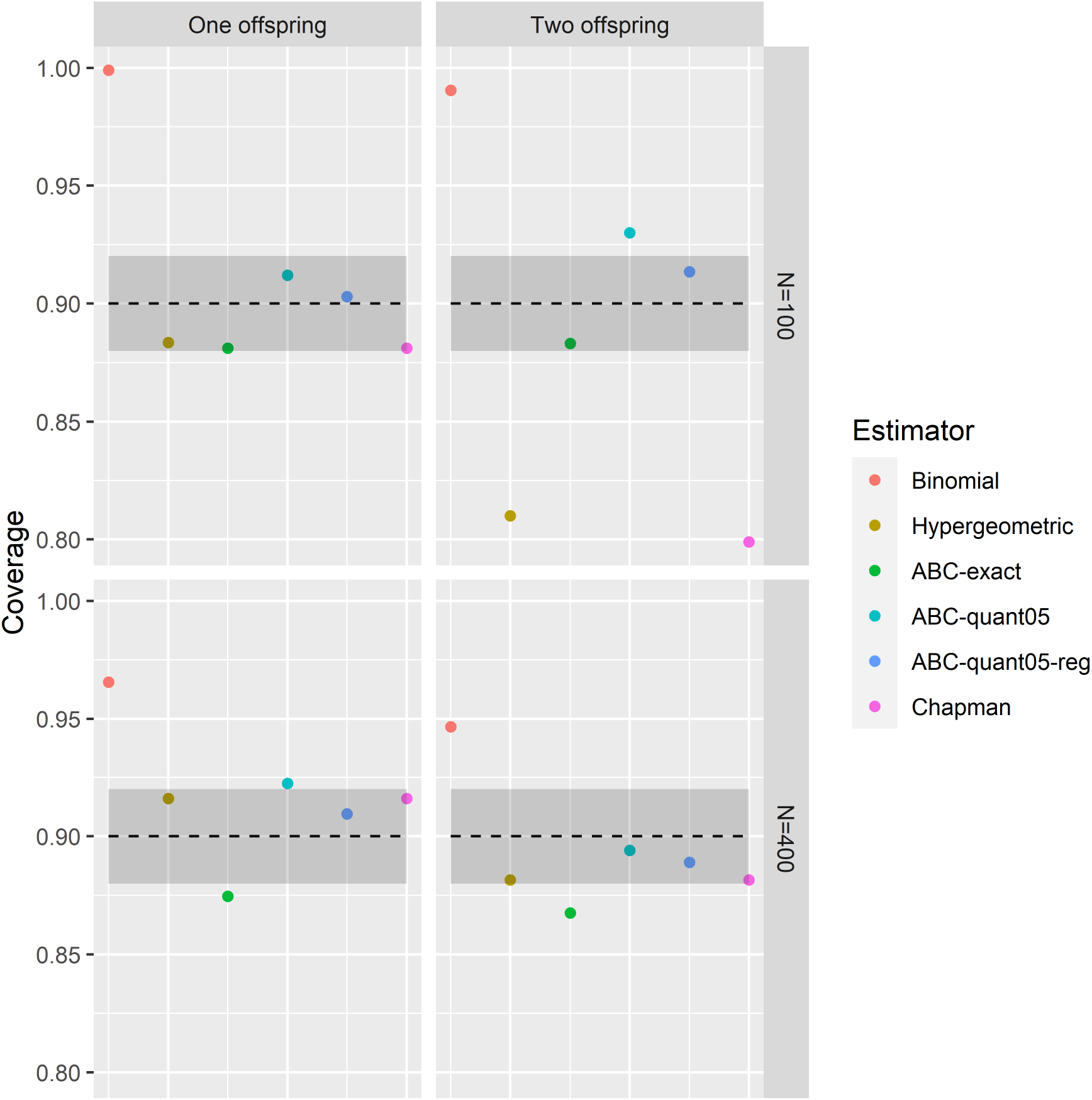
90% interval coverage for 6 estimators fitted to simulated mother-offspring close-kin mark-recapture data. I examined performance under two different adult female population sizes (100 or 400), and under scenarios where adult females had 1 or 2 offspring. The shaded area represents ± 3 SE from nominal (where SE represents Monte Carlo standard error), values outside of which are unlikely to have arisen by random chance.

### 1.2. Simulation study 2: Multiple years, population trend

In a second simulation study, I address the common situation where an investigator has access to genetic samples from a multi-year monitoring program from a species with overlapping generations. In such a case, researchers are typically interested in overall abundance as well as population trend. Here, I focus again on analysis of parent-offspring pairs (POPs), but consider an age-structured population where I assume that animal ages are known at the time of sampling (e.g., through teeth annuli or otolith rings). In addition to increased complexity, this scenario does not lend itself to a “true” likelihood (as with the hyper-geometric in the previous study), so that I only make comparisons between binomial pseudo-likelihood and ABC estimators. The number of sufficient statistics is considerably greater in this scenario, further challenging ABC methods (e.g., it will now be computationally infeasible to construct an “exact” ABC algortihm).

#### 1.2.1. Data generating model

To generate data for an age structured population, I formulated an individual-based model (IBM) where population dynamics approximated the life cycle diagram (and associated Leslie matrix, **L**) in Fig. 4. Setting *S*_0_ = 0.65, *S*_1_ = *S*_2_ = *S*_3_ = 0.9, and *f*_2_ = *f*_3_ = 1.0 leads to **L** having a dominant eigenvalue *λ*≈1.0, indicative of a stable population. I used these values to simulate population dynamics, applying the same survival probabilities to males as females, and assuming a 50/50 sex ratio at birth. Each simulation started with a population of 100 females and 100 males, the ages of which were sampled from a multinomial distribution with probabilities set to stable age proportions (the dominant eigenvector of **L**), with annual mortality of each individual simulated with a Bernoulli distribution. Following each survival interval, the number of offspring per living adult female was simulated from a Poisson distribution with expectation 2.0, where fathers of offspring were chosen randomly from the population of adult males. The population was simulated forward for 20 years, with 25 animals sampled per year in the final 10 years of the simulation. Animals were sampled randomly from the pool of newly dead animals, emulating destructive sampling or carcass surveys. I used the `fishSim` R package (Baylis, 2019) to simulate populations, conduct (virtual) sampling, and to summarize kinship of sampled individuals.

**Fig 4.**
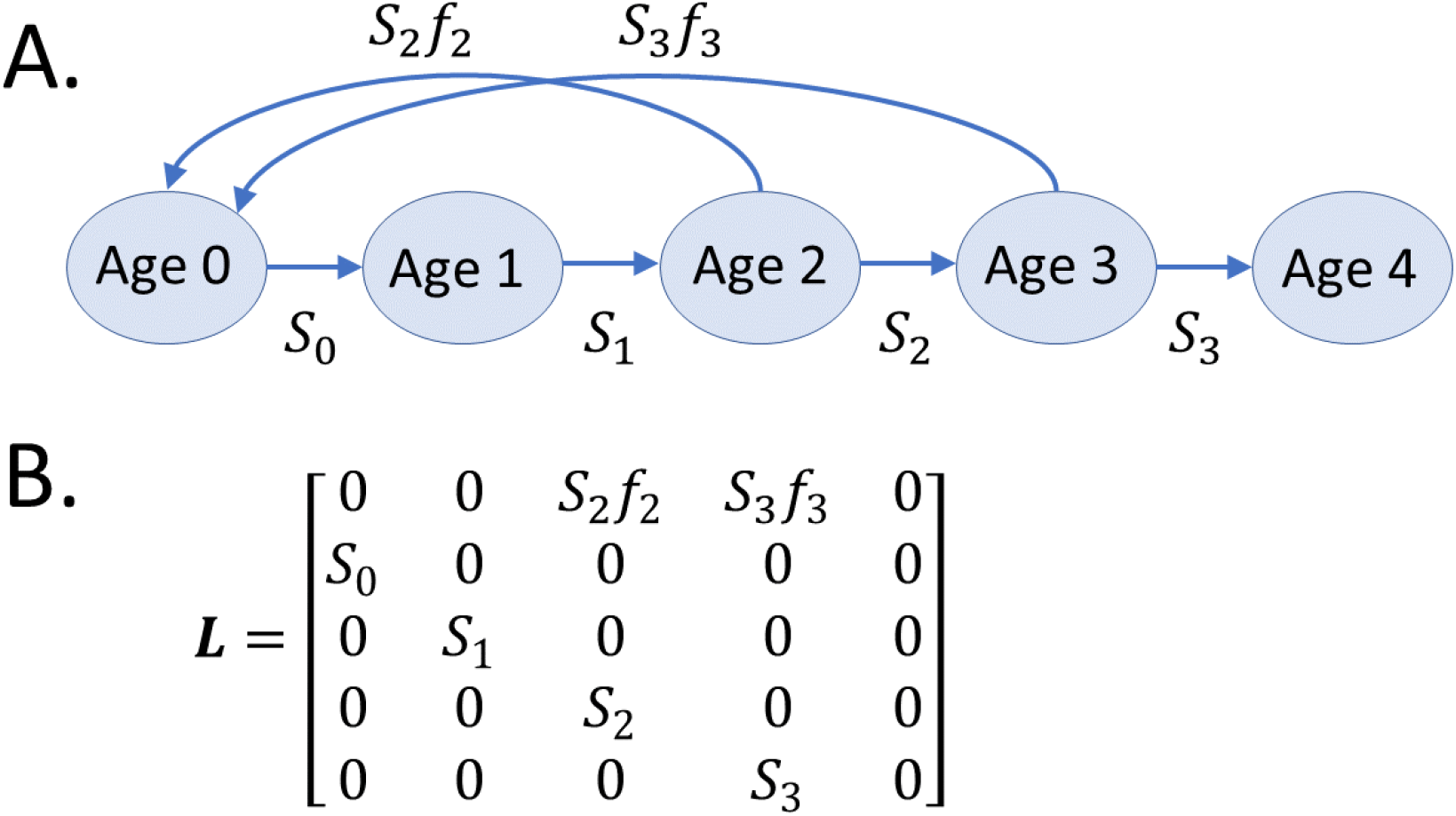
Life cycle diagram (A.) and associated Leslie matrix (B.) associated with simulation study 2. Both diagrams depict female-only dynamics with a post-breeding census, where S_a_ gives survival probability for an age a female, and f_a_ gives the expected number of female offspring per age a female. In this simplistic model, survival of 4-year-olds is set to zero.

#### 1.2.2. Estimation models

For each simulated data set, I fitted one CKMR model by minimizing a negative log pseudo-likelihood (NLPL), and another via a simulation-based ABC algorithm. Both approaches were provided true fecundity schedules and adult survival probabilities which were treated as known; estimated parameters were (i) year 1 population size, and (ii) age-0 survival. Since fecundity and the other survival parameters were fixed, first year survival effectively controlled population growth rate. However, performance (proportion relative bias, confidence/credible interval coverage) was computed relative to two derived parameters, namely *λ* and the mean number of breeders.

Sufficient statistics for the binomial pseudo-likelihood were the number of pairwise parent-offspring comparisons between individuals (*n*_*db*_) and number of successes (*y*_*db*_) summarized by year that a potential parent was sampled (*d*) and the year of birth of the potential offspring (*b*). Omitting normalizing constants, the pseudo-likelihood was product binomial, namely

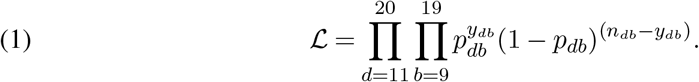

Note that I limit comparisons to those for which matches are actually possible (i.e., where the potential parent is of breeding age at the time of the offspring’s birth).

Inference about abundance and population growth comes from placing additional structure on the success probabilities, *p*_*db*_. We follow Bravington et al. (2016) in basing *p*_*db*_ on expected relative reproductive output; since males and females were assumed to have the same reproductive and survival schedules, and breeders are all equivalent in terms of reproductive value, this is simply *p*_*db*_ = 2*/B*_*b*_, where *B*_*b*_ is the number of breeders in year *b*. To compute *B*_*b*_, we rely on a forward-projecting population model, which consists of (i) an initial abundance-at-age column vector, **N**_1_ = [*N*_10_, *N*_11_, …, *N*_14_]^⊺^, and the Leslie matrix **L** in Fig. 4. As mentioned previously, all values in **L** were fixed except for *S*_0_ (age-0 survival) which is treated as a parameter to be estimated. The initial abundance-at-age vector was computed as **N**_1_ = *N*_1_***π***, where ***π*** is the dominant eigenvector of **L** (Caswell, 2001) and initial population size *N*_1_ is an additional parameter. The abundance vector in year *t, t >* 1, is then computed as **N**_*t*_ = **LN**_*t*−1_. Finally, 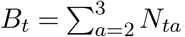, where *N*_*ta*_ gives the number of age *a* individuals in year *t*.

To get estimates of parameters, I used the ‘nlminb’ function in R (R Development Core Team, 2017) to minimize the NLPL using logit and log-links on *S*_0_ and *N*_1_, respectively. I programmed the NLPL in templated C++ code; in conjunction with Template Model Builder (TMB; Kristensen et al., 2015), this allowed for auto-differentiation and automatic computation of asymptotic standard errors for functions of parameters (e.g., *λ* and 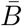). I then computed approximate log-based confidence intervals for these derived parameters as 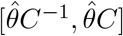, where 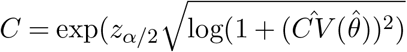, and *θ* represents a generic parameter (Buckland et al., 2001).

To implement an ABC algorithm, I relied on the same IBM that was used to simulate data. In this case, summary statistics from each IBM were the *y*_*db*_ values (number of parent-offspring matches). There were 20 possible non-zero *y*_*db*_ per IBM run; rather than suppose a particular relationship between these statistics and the parameters of interest (*λ* and 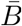), I used a neural network to relate these to **y** as suggested by Blum and François (2010) and implemented in the `abc` package in R (Csillery, Francois and Blum, 2012). Pseudo-code for the ABC algorithm is presented in Algorithm 2.

##### Algorithm 2

Approximate Bayesian computation algorithm for linear trend CKMR analysis (simulation study 2).

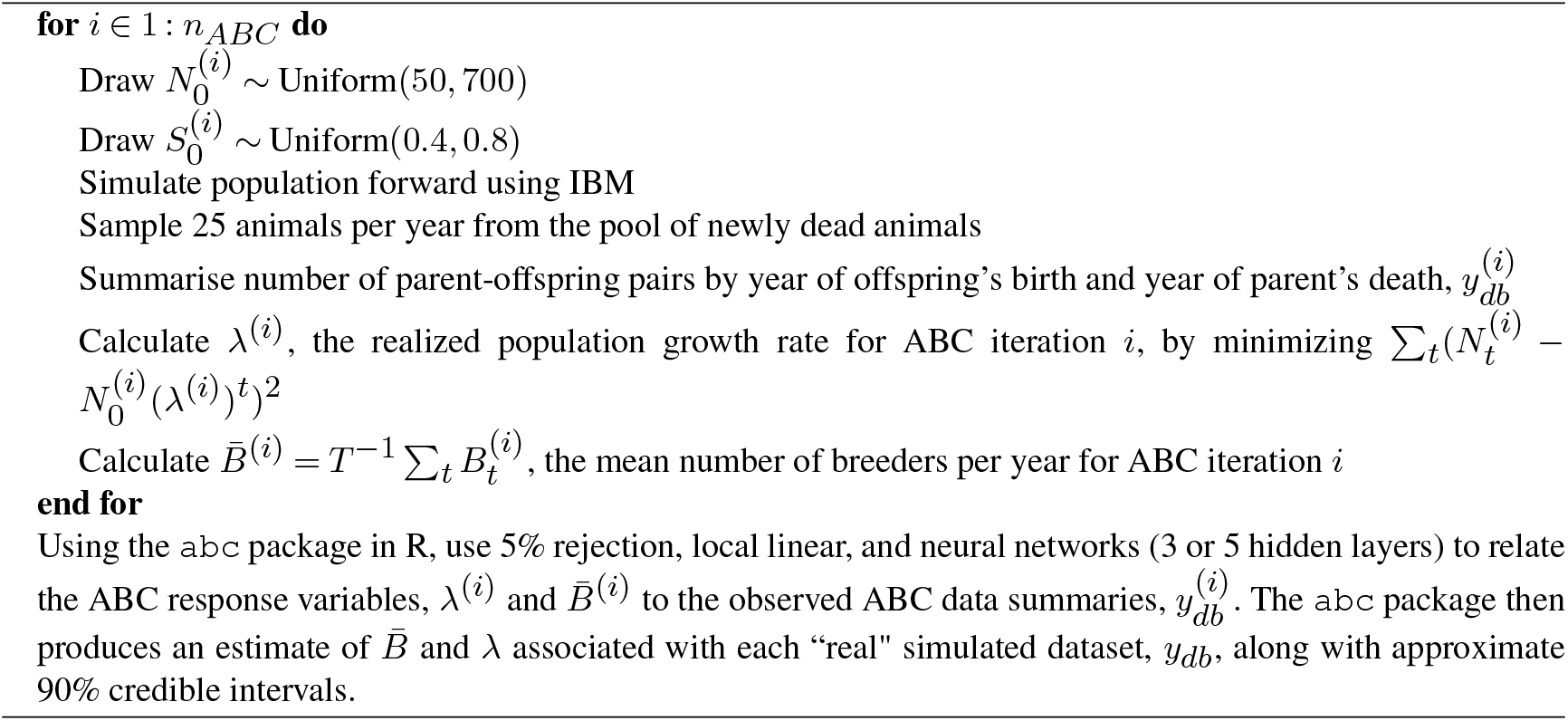

The large number of IBM model runs (18,001) performed for each simulation replicate required a substantial amount of computing power; I thus implemented the ABC algorithm in parallel. Using 12 cores on a modern PC laptop, I was able to conduct 500 simulations (a total of 9,000,500 IBM runs) in approximately 8 days. Estimator performance was judged relative to proportional relative bias and 90% confidence/credible interval coverage. For the ABC algorithms, bias was calculated relative to the posterior median, and 90% credible intervals were calculated using 5th and 95th quantiles. I employed a number of posthoc ABC algorithms from the `abc` package, including rejection sampling (5% threshold), local linear regression (Beaumont, Zhang and Balding, 2002), and neural networks (with 3 or 5 hidden layers Blum and François, 2010).

#### 1.2.3. Results

Performance differed depending on which derived parameter was of interest. If population trend (*λ*) was of interest, the ABC algorithms mostly had lower bias and mean squared error than pseudo-likelihood estimates (Fig. 5), and in two cases had credible interval coverage close to nominal. However, for the mean number of breeders, the pseudo-likelihood estimator performed better, though all estimators (ABC and pseudo-likelihood) had coverage less than nominal. All abundance estimators exhibited positive bias, with the simpler ABC algorithms resulting in high bias (10-20%). Among the ABC estimators, only the neural network with five hidden layers had a positive bias of *<* 5%.

**Fig 5.**
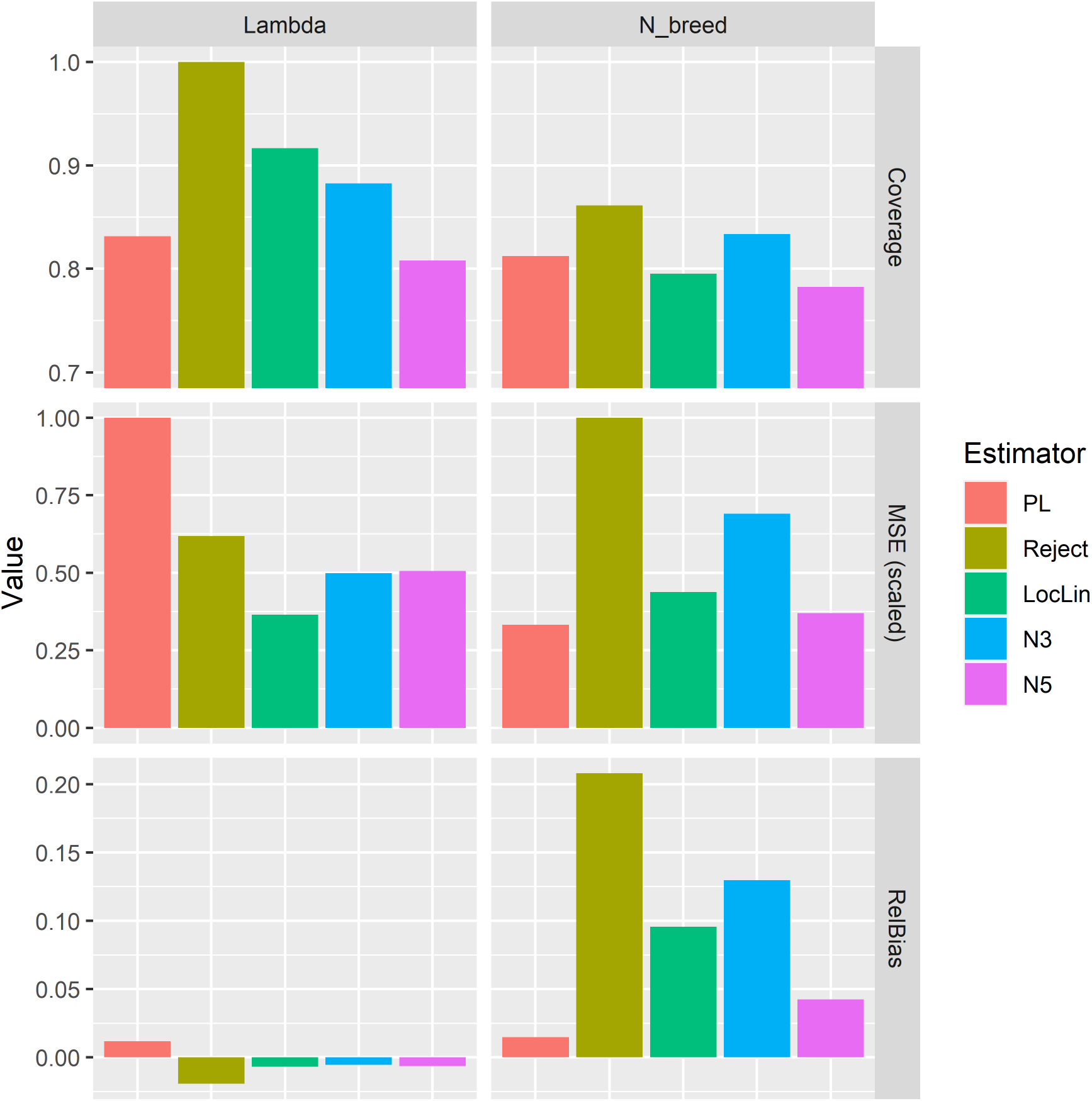
Performance of various CKMR estimators for simulation study 2. Included are 90% confidence/credible interval coverage, scaled mean squared error, and proportional relative bias for two derived parameters: finite rate of population growth (“Lambda”) and the mean number of breeders (“N_breed”). The estimators include pseudo-likelihood (“PL”), and 4 ABC estimators, including a simple rejection algorithm (“Reject”), local linear regression (“LocLin”), and neural networks with either 3 (“N3”) or 5 (“N5”) hidden layers.

### 1.3. Simulation study 3: Monogamous mating, extra-pair paternity

In a final simulation study, I examined the utility of using ABC for analyzing data where breeding is nonrandom. In particular, I developed an IBM roughly built around the demography and breeding biology of American beavers (*Castor canadensis*) which exhibit monogamous mating. Demographic processes were again specified through an age-structured discrete-population model with a post-breeding census and annual time steps (similar to Fig. 4), with a maximum age of 15. Annual survival probabilities were set to *S*_0_ = 0.39 (kit survival); *S*_1_ = *S*_2_ = 0.65; *S*_3_ = *S*_4_ = … = *S*_13_ = 0.77; *S*_14_ = 0. Values of juvenile (ages 1-2) and adult survival (ages 3+) were informed by the literature (McNew Jr and Woolf, 2005; Smith et al., 2016), whereas kit survival was adjusted so as to achieve roughly stable populations. Beavers were assumed sexually mature starting at age 3, and the number of offspring per year for partnered adult females was drawn from a Poisson distribution with expectation 4.0.

To implement a monogamous IBM, I started with some of the core functions from the `fishSim` packages, but created a new mating function. At the start of each year, each unpartnered sexually mature female was randomly assigned to an unpartnered sexually mature male; unpartnered females did not reproduce. IBM simulations were initialized with a population size of 200 (initial ages were sampled from a multinomial distribution with success probabilities set to stable age distribution proportions from the associated Leslie matrix model). Each IBM simulated mortality and birthing events for 30 annual time steps (2 generations), with virtual genetic samples obtained from 20 newly dead animals in each of the last 15 years.

The monogamous mating structure means that full thiatic pairs (FTPs; e.g, full aunt-niece) are expected to be common. In pseudo-likelihood analyses, their presence would typically preempt use of HSPs for CKMR inference, given that FTPs have the same proportion of co-inherited alleles as HSPs and GGPs. In absence of FTPs, GGPs and HSPs could be modeled with a mixture distribution (e.g., Bradford et al., 2018; Taras et al., 2024), or HSP analysis could simply be limited to comparisons where prospective birth intervals are less than 2 times the age of sexual maturity. However, to our knowledge no one has written a probabilistic model for FTPs, and their birth interval distributions considerably overlap HSPs (see, e.g., Fig. 6). Thus, pseudo-likelihood inference would likely be limited to POPs, or perhaps POPs and full sibling pairs (FSPs; see e.g. Patterson et al., 2022).

**Fig 6.**
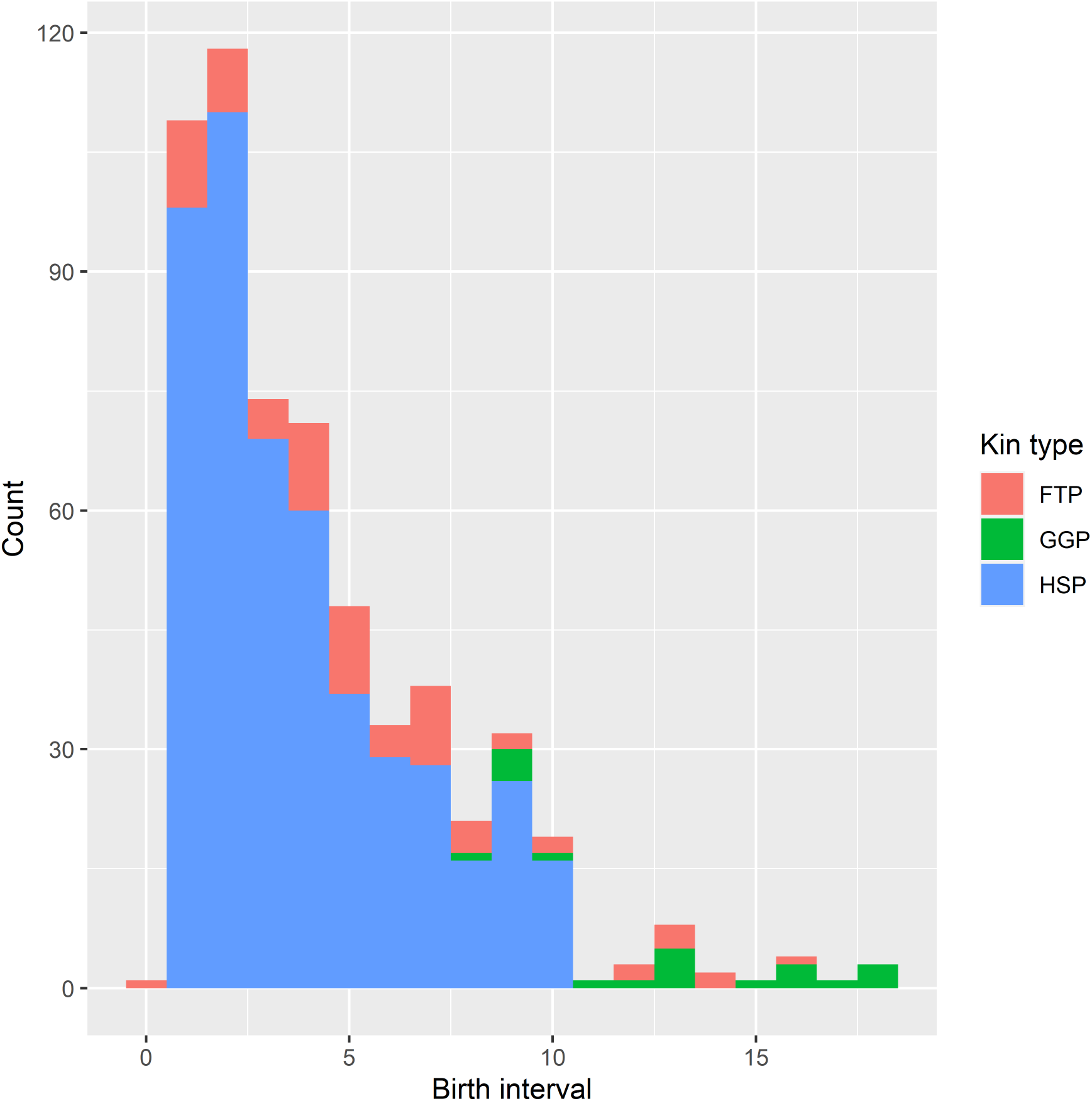
Frequency of half-sibling (HSP), full-thiatic (FTP) and grandparent-grandchild (GGP) pairs obtained from simulated genetic sampling applied to a single IBM with monogamous mating. These kin types have the same expected proportion of coinherited alleles and cannot be distinguished using most modern genotyping methods.

A possible alternative is to use the ABC machinery to include all HSP+ matches (HSP+FTP+GGP) as additional data without needing to explicitly supply the probability of a match. That is the approach I take in this simulation study, where I employ POPs, FSPs, and HSP+ matches for inference. To identify appropriate statistics to use for the ABC algorithm, it is worth considering the probabilities often associated with various types of kin pairs. For instance, the probability of a half-sibling match is often constructed as proportional to

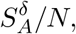

where *S*_*A*_ is adult survival, *δ* is the difference in birth years of the potential HSPs being compared, and *N* is abundance (see, e.g., Bravington et al., 2016). In essence, the birth gaps of siblings carry information about adult survival probabilities. By contrast, the probability of parent-offspring pairs does not include a survival term. From these observations, I propose to use frequencies of FSP and HSP+ as summary statistics for survival, and the frequencies of all kin pair types (POP, FSP, and HSP+) as summary statistics for abundance. Specifically, for survival I use *y*_Δ*t,k*_, the number of pairs of type *k* found that have differences in birth interval of Δ_*t*_ = 1, 2, · · ·. For FSP and HSP+ summary statistics for abundance and trend, I use 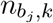 : the number of kin pairs for which the older animal had birth year *b*_*i*_. For POPs for abundance and trend, I use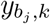, the number of kin pairs for which the potential offspring (i.e., younger animal) has birth year *b*_*j*_. Eliminating summary statistics that had zero variance, there were a total of 21 summary statistics for survival, and 49 summary statistics for abundance and trend.

Pseudo-likelihood approaches to inference would also make use of the number of comparisons as a binomial index (e.g.,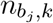), but in the IBM setting we can control for sample size by requiring the number of tissue samples in ABC replicates to be equal to that observed in the “real data” (in our case, 20 samples per year for the final 15 years of simulation). Of course, some ABC replicates, particularly those with small or declining populations, might not be able to meet this level of sampling. These cases were automatically rejected when conducting ABC post-processing.

Overall, the simulation study was structured as follows:

- For a total of 200 simulation replicates, mating and population dynamics were simulated using an IBM, with sampling as described above;
- For each simulation, a total of 24,000 ABC replicates were generated, each using the same IBM model as used to generate the “real” data set. However, each ABC replicate used the following iid draws:

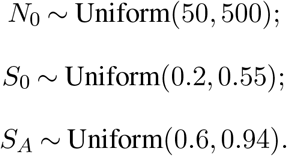
- Summary statistics were recorded for each ABC iteration where the required amount of tissue samples were available to be collected;
- For each simulation, 2 ABC algorithms were applied: one using a 5-layer neural network within the `abc` package, and one using a semi-automated algorithm similar to that described by Fearnhead and Prangle (2012); see Algorithm 3. For the latter, I set *q* = 0.02.

I used standard performance metrics (relative bias, credible interval coverage, RMSE) to gauge estimator performance. These were computed relative to posterior predictive modes, as approximated with kernel density estimators applied to ABC output. Performance was assessed for three quantities: the realized finite rate of population growth (*λ*), the mean abundance of female breeders (*N*_*f*_), and adult survival probability (*S*_*A*_).

#### 1.3.1. Results

I summarized results from 158 successful simulations, with the remaining 42 being censored because demographic stochasticity prevented pre-allocated sample sizes (20 newly dead females per year) from being collected. Posterior modes for ABC inference had negligible bias for population growth rate (*λ*) and adult survival (*S*_*A*_), but exhibited positive bias (e.g. 15%) for breeding female abundance (Table 1). Performance as a whole was better for the semi-automated algorithm than the neural network approach, which had credible intervals that were considerably too short.

**Table 1.**
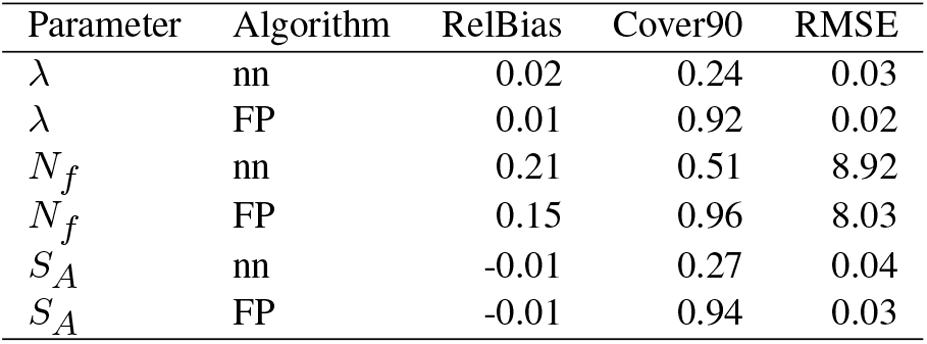
Performance of two ABC algorithms when estimating the finite rate of population change (λ), the abundance of adult females (N_f_), and adult survival probability (S_A_) from simulated CKMR data from American beaver populations exhibiting monogamous mating behavior. The two ABC algorithms included a neural network (“nn”) implemented in the `abc` R package (Csillery, Francois and Blum, 2012), and a semi-automated algorithm (Fearnhead and Prangle, 2012) (“FP”). Performance metrics include proportion relative bias (“RelBias”), 90% credible interval coverage (“Cover90”), and root mean squared error (RMSE). Bias and RMSE were computed relative to the posterior mode, and credible intervals were constructed using 5th and 95th quantiles.

## 2. Example: Caribou data

I next applied simulation-based inference to estimate the population size and trend of the Tonquin population of boreal caribou (*Rangifer tarandus*) in Alberta, Canada (McFarlane et al., 2018; Merriell, Manseau and Wilson, 2024). Data from this population were recently used to examine the performance of pseudolikelihood CKMR estimates of abundance (Merriell, Manseau and Wilson, 2024). Such estimates can be compared to more longitudinal estimates of abundance produced with more traditional capturemark-recapture (CMR) estimates. In particular, Merriell, Manseau and Wilson (2024) found that CKMR estimates of abundance compared favorably with CMR estimates, although confidence intervals were considerably broader for the former.

Although the caribou data are thoroughly documented elsewhere (e.g., McFarlane et al., 2018; Merriell, Manseau and Wilson, 2024), some details of the study population and sampling program are worth describing here. First, birthing of caribou calves happens in the spring (May to mid-June), and DNA was obtained during the winters of 2006-2015 by collecting fecal pellets. The quality of the DNA was sufficient for determining parent-offspring relationships, but no further (e.g., no half-siblings). Hormones in fecal pellets were used to distinguish calves (age ≈6 months) from non-calves. However, female sexual maturity first reliably occurs at age 3, so that the non-calf class includes a limited number of age 1.5 and 2.5 non-breeders. Each pregnant female gives birth to one calf.

### Algorithm 3

Semi-automated approximate Bayesian computation algorithm for multi-year close-kin mark-recapture data

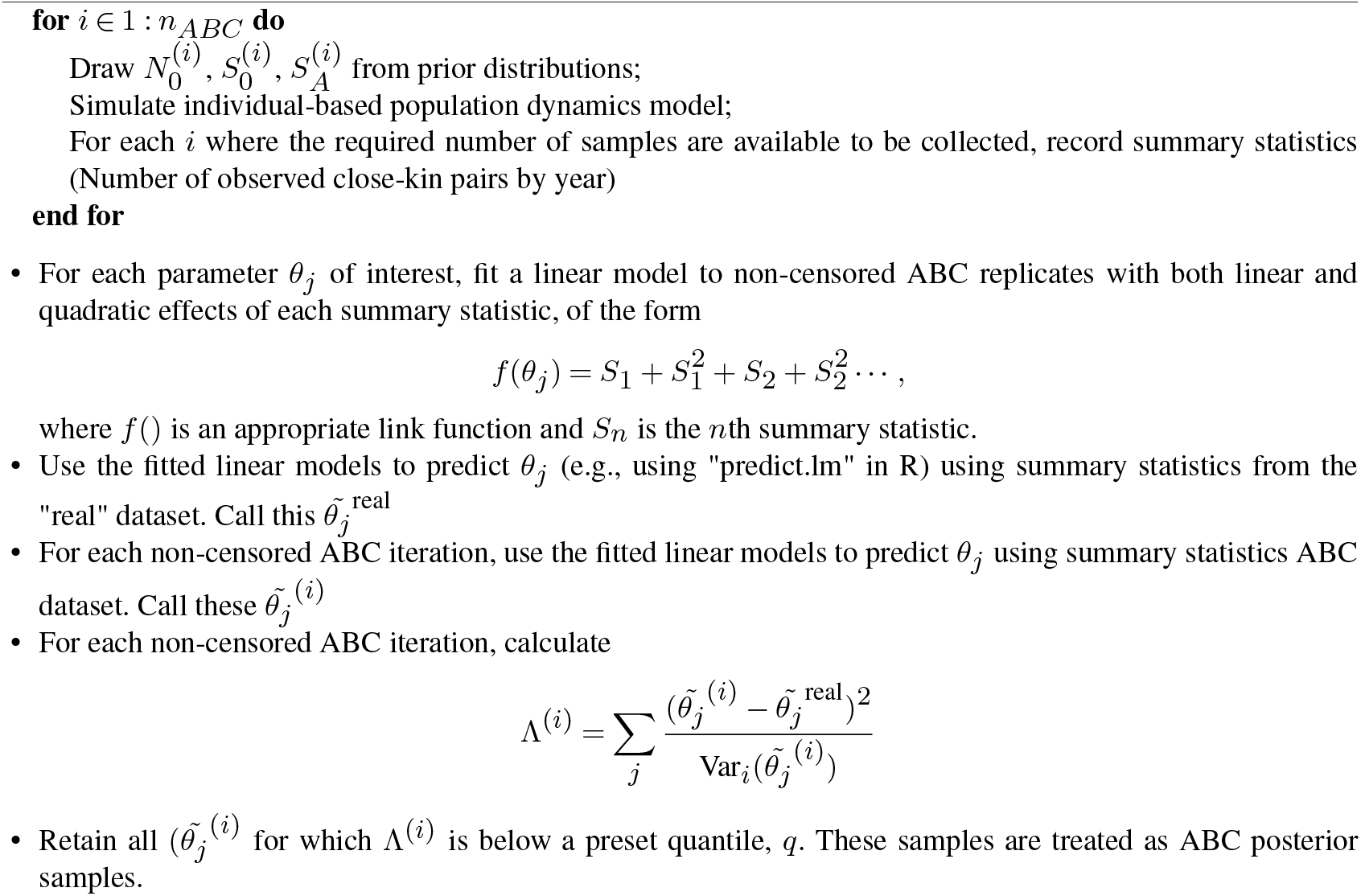

I used the reported number of sampled caribou, together with the number of observed mother-offspring pairs in each year to fit several CKMR models. The first simply used the Chapman estimator (see section 1.1.2) and logit-transformed intervals (Sadinle, 2009) applied to data from individual years. These are compared to the CMR estimates from McFarlane et al. (2018) and pseudo-likelihood based CKMR estimators from Merriell, Manseau and Wilson (2024) (Fig. 7). In general the three methods provided similar estimates, though the Chapman estimator tended to be lower than the pseudo-likelihood estimator. The Chapman estimator also had considerably shorter confidence intervals, as might be expected given the previously noted tendency of pseudo-likelihood estimators to be overly conservative when applied to small populations.

**Fig 7.**
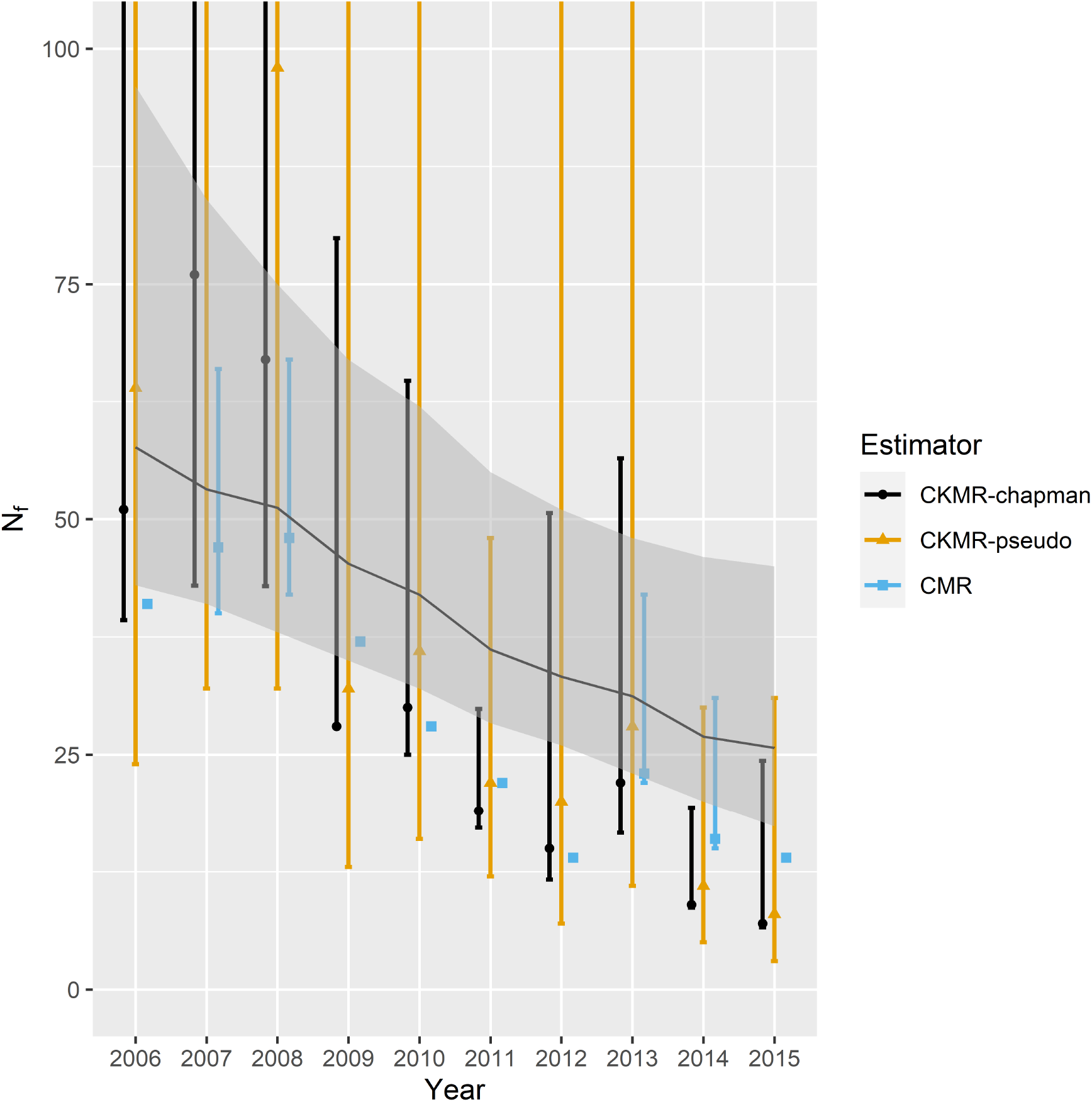
A comparison of female abundance estimators for the Tonquin caribou population, 2006-2015. Points and vertical bars represent single-year abundance estimators and 95% intervals, including standard capturemark-recapture (“CMR”; taken from McFarlane et al., 2018); a CKMR estimator based on pseudo-likelihood (“CKMR-pseudo”; taken from Merriell, Manseau and Wilson, 2024); and a Chapman CKMR estimator. Also shown is a multi-year, simulation-based ABC posterior mode with 95% credible intervals (thin black line with light gray envelope).

In addition, I used a semi-automated ABC algorithm (sensu Fearnhead and Prangle, 2012) to estimate parameters of a multi-year IBM fitted to all of the CKMR data (Algorithm 3). A total of 1.2 million IBM runs were performed, each consisting of birth, death, and pellet collection processes occurring over 10 years with virtual caribou having a maximum age of 20. I set yearling and subadult (ages 1-2) survival probability to 0.8 and adult (age 3+) survival probability to 0.85, values that were used in Merriell, Manseau and Wilson (2024). Similarly, I set the expected number of offspring to zero for calves and yearlings, to 0.15 for age 2 females, and to 0.9 for age 3+ females (Merriell, Manseau and Wilson, 2024). These values were treated as fixed for all IBM simulations. Variability in abundance and trend was induced by initializing each population with a different number of initial individuals (*N*_0_ ∼ Uniform(50, 400)) and calf survival probability (*S*_0_ ∼ Uniform(0.05, 0.65)). Note that asymptotic population growth rate (*λ*) is then a one-to-one, monotonic function of *S*_0_. The initial age and sex structure of each IBM was set by simulating from a multinomial distribution with cell probabilities set to the asymptotic age distribution determined from the associated Leslie matrix (Caswell, 2001), and assuming a 50/50 sex ratio. The model was implemented with a 6 month time step; I simulated DNA collection from fecal pellets in the winter and births in the spring, using the same number of sampled calves and potential mothers as in the real caribou dataset. Summary statistics were the number of female-calf pairs in each of the 10 winters. For reference, the real caribou dataset (Merriell, Manseau and Wilson, 2024, Table 5) consisted of 3-11 calves sampled per year, 8-32 potential mothers sampled per year, and 1-8 mother-offspring pairs per year. The semi-automated ABC algorithm used a quantile of *q* = 0.002.

In general, multi-year ABC estimates of female abundance showed the same population trend as the single year estimators (Fig. 7). However, on average absolute abundance was higher than the CMR and Chapman CKMR estimators. This may reflect the fact that it is difficult to sustain the observed level of pellet collection samples in small populations. Indeed, a large proportion of simulations with small population sizes were censored for this reason (see e.g. Fig. 8). Ungulate populations often exhibit stochasticity in annual recruitment, so that greater numbers of calves would be available to be sampled in high recruitment years. By contrast, a simple two parameter IBM, assuming constant recruitment, may only allow higher levels of sample collection when populations on the whole are more abundant.

**Fig 8.**
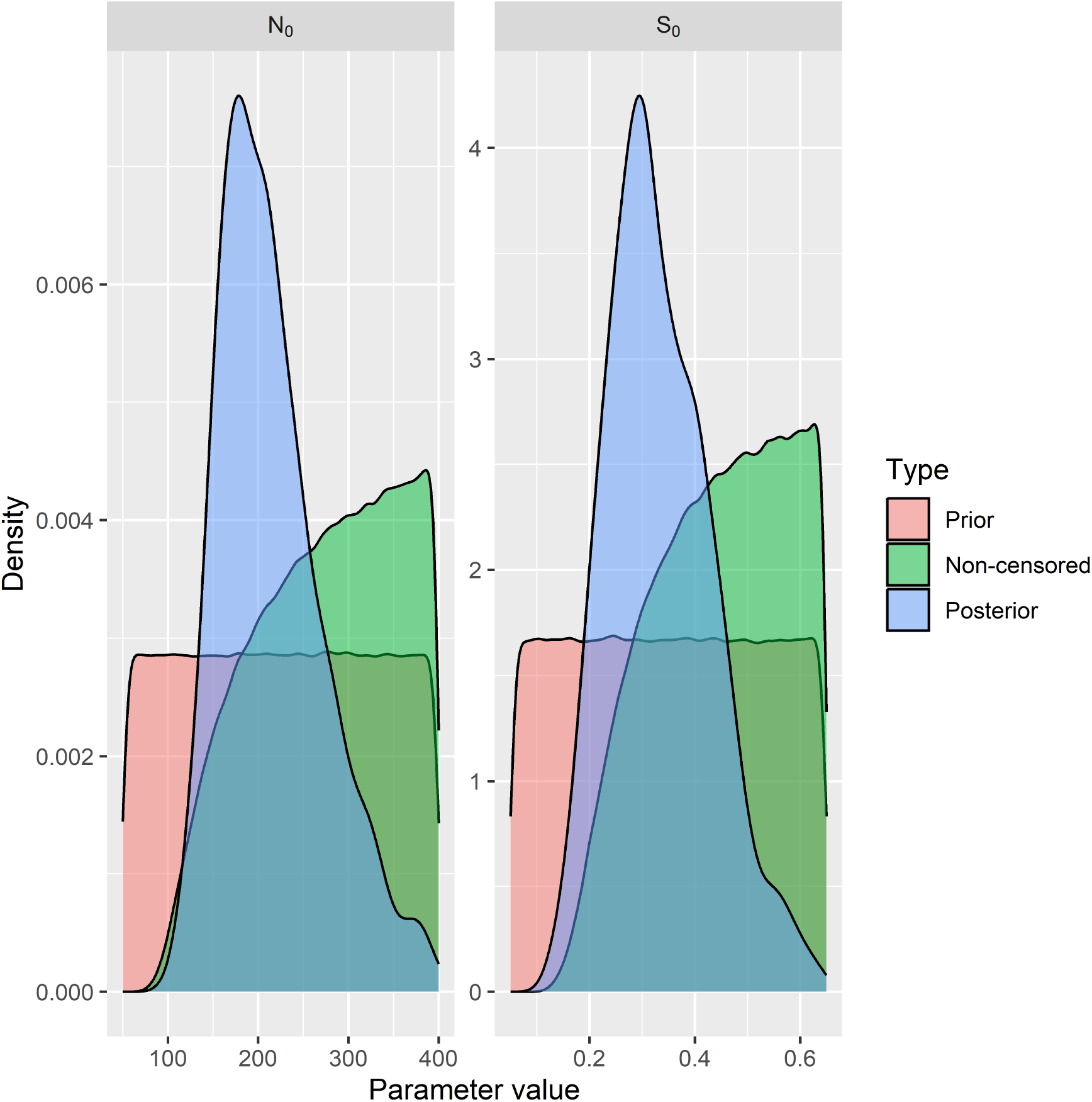
Prior and posterior distributions of parameters when conducting ABC-based inference from multi-year caribou mother-offspring CKMR data. The parameters are N_0_, the initial population size at the start of simulation time, and S_0_, calf survival. Many of the ABC simulation replicates with low population size or low calf survival were censored because they drove the population too low to sustain the number of fecal pellet samples achieved in the real data set.

## 3. Discussion

Close-kin mark recapture is in its infancy, with most applications to-date using a binomial pseudo-likelihood (Bravington et al., 2016) for inference. This formulation works well for large populations and species with well behaved life histories, but can result in understated precision when applied to small populations (Merriell, Manseau and Wilson, 2024). The pseudo-likelihood approach also requires specifying analytical formulae for the probabilities of kin pair matches, which can become complicated when populations are spatially structured (Conn et al., 2020; Davies, Bravington and Thomson, 2017), feature skip breeding or other reproductive inconveniences (Sévêque et al., 2024; Swenson et al., 2024), or when aging errors occur (Swenson et al., 2024). On the extreme end, pseudo-likelihood probabilities are potentially intractable, as in monogamous mating structures where one cannot reliably discriminate between HSP, GGP, and FTP relationships.

In this paper, we confirmed the results of Merriell, Manseau and Wilson (2024) that pseudo-likelihood inference results in relatively unbiased but conservative estimates when applied to small populations, in the sense that confidence intervals are far too wide. By contrast, alternative estimators performed better, including (1) a Chapman estimator applied to single-season CKMR data, and (2) simulation-based estimators using ABC. The Chapman estimator may only be applied in a very limited circumstance (single season data, one offspring per female), so simulation-based inference is the most versatile alternative. We demonstrated reasonable performance of ABC estimators in several simulation studies, including multi-year studies. Performance was best for survival and population trend parameters; absolute abundance estimates tended to be positively biased.

Application of simulation-based inference seemed to be somewhat sensitive to the particular ABC algorithm that was employed. For relatively simple problems (as with single year mother-offspring data), simple rejection-based ABC algorithms may suffice. For others, it seems necessary to expend effort tailoring the algorithm to the data at hand. For instance, in simulation study 3, a bespoke semi-automated algorithm (sensu Fearnhead and Prangle, 2012) seemed to perform better than a neural network algorithm available in the `abc` R package (Blum and François, 2010; Csillery, Francois and Blum, 2012). This comparison is not meant as a criticism of `abc`. I suspect this result is circumstance specific, and cannot rule out the possibility that someone intimately familiar with the package could have achieved more favorable results. Regardless, knowledge of the sufficient statistics needed for pseudo-likelihood inference can lead to intuition about what statistics carry information about different parameters, and thus for setting the maximal number of summary statistics for different parameters needed for ABC inference.

One challenge with applying ABC methods to small populations is the high probability of censoring ABC replicates (i.e., IBM runs) with low abundances or high population declines. Often in these situations, the population can drop below the level needed to sustain the level of sampling in the observed data. I suspect that this is the primary reason for negative bias observed in multi-year abundance estimates. This phenomenon also seems related to model misspecification - for instance, a simple 2- or 3-parameter IBM may require a comparably large population size to meet required sample sizes, whereas an IBM with more parameters reflecting annual changes in recruitment or survival might be able to meet sample size requirements with a lower population size. An obvious alternative would be to include a larger suite of parameters within ABC inference; however, the types of ABC algorithms explored in this paper are subject to the curse of dimensionality (Beaumont, 2010), and can only reliably handle a small number of parameters. Although these are attractive because IBMs can be run in parallel, a larger parameter space may necessitate a sequential algorithm such as ABC-MCMC (Marjoram et al., 2003), or perhaps synthetic likelihood (Wood, 2010).

The simulation studies used in this paper were computationally demanding, in the sense that each simulation requires a large number of ABC replicates. As a consequence, I reduced both the total number of simulations and the total number of ABC replicates per simulation to values lower than typically used in the literature. For instance, simulation study 3 consisted of 200 simulations, with 24,000 ABC samples per simulation. By contrast, many studies employing ABC use millions of ABC samples when analyzing a given data set. In general, I suspect that performance, both in terms of bias and coverage, will improve relative to the results presented here when using a higher sample size. That said, subsetting the existing simulations to a smaller number (e.g. 24,000 to 12,000 in simulation study 3) did not in itself noticeably reduce performance.

Although the simplicity of simulation-based inference is attractive, there are potentially more life history values that need to be fixed by an analyst than is typically required for CKMR estimators. For instance, when setting up IBMs for this paper, I effectively modeled all age classes, fixed reproductive rates and some survival probabilities, and dictated how (and with whom) individuals were allowed to mate. Since CKMR does not provide information about pre-breeding survival, this requires using auxiliary information to set many such values. Similarly, reproductive schedules were assumed known. Thus, in implementing simulation-based CKMR models, one may be opening oneself up to more assumption violations than with pseudo-likelihood estimators. The quality of simulation-based inference will thus likely be predicated on having high quality life history data for the population at hand.

Despite these potential methodological and assumption-related hurdles, simulation-based inference holds great promise for CKMR inference, particularly for small populations. Many processes can potentially be integrated over, including spatial structure, aging error, and many of the other real life complications that make specifying pseudo-likelihood probabilities difficult. It is comparably much easier to construct an IBM that encapsulates the main features of population dynamics and to compare resulting discrepancy measures to those arising from a real life dataset. This potentially opens the door for more widespread use of CKMR methods for assessing the abundance and demography of animal populations.

## Acknowledgments

I thank J. Ver Hoef and B. Brost for providing valuable feedback on a previous version of this manuscript. Code and data for this study are currently available on github at https://github.com/pconn/CKMR_ABC/tree/master and will be archived to a permanent repository following manuscript acceptance.

